# Classifying Mild Cognitive Impairment from Normal Cognition: fMRI Complexity Matches Tau PET Performance

**DOI:** 10.1101/2025.01.16.633407

**Authors:** Kay Jann, Gilsoon Park, Hosung Kim, the Alzheimer’s Disease Neuroimaging Initiative

**Author notes:** Corresponding author Address: Kay Jann, PhD, Mark and Mary Stevens Neuroimaging and Informatics Institute Keck School of Medicine, University of Southern California 2025 Zonal Ave, Los Angeles, CA, 90033. Data used in preparation of this article were obtained from the Alzheimer’s Disease Neuroimaging Initiative (ADNI) database (adni.loni.usc.edu). As such, the investigators within the ADNI contributed to the design and implementation of ADNI and/or provided data but did not participate in analysis or writing of this report. A complete listing of ADNI investigators can be found at: http://adni.loni.usc.edu/wp-content/uploads/how_to_apply/ADNI_Acknowledgement_List.pdf. Equal first author contribution (Kay Jann, Gilsoon Park).

## Abstract

**Background:** Tau protein accumulation is closely linked to synaptic and neuronal loss in Alzheimer’s disease (AD), resulting in progressive cognitive decline. Although tau-PET imaging is a direct biomarker of tau pathology, it is costly, carries radiation risks, and is not widely accessible. Resting-state functional MRI (rs-fMRI) complexity—an entropy-based measure of BOLD signal variation—has been proposed as a non-invasive surrogate biomarker of early neuronal dysfunction associated with tau pathology.

**Objectives:** To determine whether fMRI-based brain complexity (sample entropy and multiscale entropy) can match or exceed tau-PET in classifying cognitively normal (CN) versus cognitively impaired (MCI/AD) individuals. And to investigate and compare the most influential network regions-of-interest (ROIs) for classification between fMRI complexity and tau-PET, thereby identifying key neuroanatomical correlates of AD-related changes.

**Design:** A cross-sectional study employing 3D convolutional neural network (CNN) classification with five-fold cross-validation and leave-one-network-out analysis.

**Setting:** Data from the Alzheimer’s Disease Neuroimaging Initiative (ADNI) database.

**Participants:** One hundred forty-seven older adults (age 72.5 ± 7.5 years), including 95 CN, 45 MCI, and 7 AD.

**Measurements:** We created whole-brain complexity maps from rs-fMRI and standardized uptake value ratio (SUVR) maps from tau-PET. Each modality was separately fed into CNN classifiers. Region-based analyses (leave-one-network-out) were performed to identify critical ROIs for classification.

**Results:** fMRI complexity showed classification accuracy comparable to tau-PET yet surpassed it in F1-score (0.64 vs. 0.61) and area under the curve (AUC; 0.73 vs. 0.67). Salience and dorsal attention networks contributed most to fMRI-based classification, and a dorsal attention network contributed most to tau-PET-based classification.

**Conclusions:** fMRI complexity performs similarly to tau-PET in detecting cognitive impairment related to AD and identifies partially distinct critical ROIs, suggesting an alternative, radiation- free imaging biomarker for earlier detection and broader clinical application.

## 1. Introduction

Cognitive decline in older age is a hallmark diagnostic criterion in the progression of dementia and Alzheimer’s disease (AD). Unfortunately, by the time AD is diagnosed, structural brain changes are often irreversible, making treatment less effective. As a result, research is increasingly focused on identifying early biomarkers, especially during the mild stages of cognitive decline, to enable earlier intervention and more effective treatment strategies.

Positron Emission Tomography (PET) and Magnetic Resonance Imaging (MRI) are the foremost imaging modalities used to identify neurophysiological and brain structural alterations in the progression of dementias. In the pursuit of classifying AD through advanced machine learning approaches, Amyloid PET and Tau PET imaging have highlighted brain areas crucial for diagnostic precision. Evidence shows that amyloid beta (Aβ) can accumulate in brain tissue 10-15 years before symptoms appear.[1,2] While amyloid deposition is seen in pre-symptomatic AD, tau accumulation correlates more closely with cognitive decline, making Tau PET a more direct biomarker of neuronal injury and subsequent cognitive impairment.[3] However, PET imaging has radiation exposure risks, is costly and not available everywhere.

Recently, brain entropy mapping based on resting-state functional MRI (rs-fMRI) has been proposed to identify abnormal brain function linked to cognitive decline.[4–6] Brain entropy estimates the complexity of Blood Oxygen Level Dependent (BOLD) signals and is hypothesized to reflect the brain’s capacity of information processing and cognitive flexibility. As cognitive functions deteriorate, BOLD signal complexity decreases, particularly in areas like the hippocampus and default mode network, which are critical for memory. This reduction in complexity is especially pronounced in AD.[7] Importantly, fMRI-complexity is very sensitive to early, subtle cognitive changes, thus offering insights into early stages of AD[8] and potentially serving as a biomarkers of mild cognitive impairment (MCI) in AD progression. Furthermore, we recently demonstrated that fMRI-complexity is negatively correlated with tau-PET uptake ratios in specific brain areas within the temporal, parietal and prefrontal cortices,[6] establishing a relevant pathophysiological relationship between these two imaging modalities.

Classifying individuals with MCI using neuroimaging markers has been challenging due to the subtlety and heterogeneity of early AD progression. This study aimed to evaluate the ability of fMRI complexity to distinguish MCI/AD from cognitively normal subjects using a deep learning classifier. We also compared the classification performance of fMRI complexity with tau-PET to assess whether fMRI complexity could serve as a surrogate marker for early AD. Finally, we investigated the specific brain regions that contributed most to accurate classification.

## 2. Methods

### 2.1 Study population

To train the classifier we used data obtained from the Alzheimer’s Disease Neuroimaging Initiative (ADNI) database (http://adni.loni.usc.edu). The ADNI was launched in 2003 as a public-private partnership, led by Principal Investigator M.W. Weiner, MD. The primary goal of ADNI has been to test whether serial MRI, PET, other biological markers, and clinical and neuropsychological assessment can be combined to measure the progression of MCI and early AD. We identified ADNI participants that had complete imaging sets for tau-PET (tracer: 18F- AV1451) one fMRI scan and a T1 weighted anatomical scan. The identified dataset consisted of 147 subjects (age=72.5±7.5, male/female = 60/87, Cognitive Normal (CN)/MCI/AD = 95/45/7). Details of clinical diagnosis in ADNI have been described in [9]: CN participants had a Mini-Mental State Examination (MMSE) score of < 26 and a Clinical Dementia Rating (CDR) of 0. Participants with probable AD had an MMSE score < 24, and a CDR of 0.5 or 1.0. MCI, early MCI, and late MCI were pooled into one MCI group.

### 2.2 fMRI data acquisition and processing

fMRI data acquisition in the ADNI basic protocol used gradient-echo echo-planar imaging in PA phase encoding direction at an isotropic voxel size of 3.4×3.4×3.4 mm^3^, with ms, for a total of 197 volumes resulting in an acquisition time or 10 Date underwent standard preprocessing including regression of physiological and motion related signal fluctuations from eroded white matter and cerebrospinal fluid masks, generated from probabilistic tissue segmentation masks of T1w-images, and 12 motion parameters (x,y,z translation and rotation plus first derivatives), co-registration to T1w structural images, normalization to MNI standard space and spatial smoothing with an 8mm Full Width at Half Maximum (FWHM) Gaussian kernel.

fMRI-complexity maps were computed based on sample entropy at six temporal scales using pattern matching lengths of m=2 and pattern matching threshold of r=0.5. We used our in-house software for this analysis step (LOFT Complexity Toolbox; github.com/kayjann/complexity). Finally, multi-scale entropy maps were created by averaging complexity maps across all temporal scales.

### 2.3 PET data acquisition and processing

Raw dynamic Tau-PET data were averaged across time, co-registered to individual T1 weighted anatomical scans and smoothed with an 8mm FWHM Gaussian Kernel. Parametric Standardized uptake value ratio (SUVR) values were then computed using a reference region in the inferior cerebellum using SUIT (https://www.diedrichsenlab.org/imaging/suit.htm).[10] Finally, tau-PET SUVR maps were normalized into MNI space.

### 2.4 Creation of Regions of Interest and Sub-Networks

To assess the performance of distinct neuroimaging approaches in classifying CN and MCI/AD, we first used a gray matter (GM) probability map derived from SPM and binarized it to mask the Sample Entropy (SampEn), Multiscale Entropy (MSE), and tau-PET SUVR maps within the GM region. This initial masking procedure ensured a consistent focus on the cortical GM region for comparison across different imaging approaches. After this process, 9 functional network regions were generated to assess the impact of specific region-of-interest (ROI)s on classification performance. These regions were based on an extended version of the 7 networks (sensorimotor, frontoparietal, dorsal attention, ventral attention with language, default mode, visual, and limbic) defined by Yeo, Krienen, Sepulcre*, et al.* [11], by additionally defining two more regions for salience and auditory networks. The networks were defined on the cortical GM in the MNI space using the Automated Anatomical Labeling (AAL) Atlas. By registering individual images into the MNI template, these 9 network ROIs were mapped on each individual image. We then conducted a systematic assessment of the contribution of each network region to the classification through a leave-one ROI-out strategy, which is based on the occlusion sensitivity analysis in the field of computer vision (see 2.6). [12,13]

### 2.5 3D Convolutional Neural Networks for Classification

To evaluate the ability of fMRI-complexity and tau-PET images to predict cognitive impairment in relation to AD progression, 3D convolutional neural networks (CNN) were used to classify individuals into CN or MCI/AD groups (Fig. 1). A cortical GM mask was applied to all images to isolate and highlight the cortical GM region as the brain region of interest. We did not apply an additional intensity normalization to utilize the original intensity values of complexity and SUVR in each image. Each of fMRI complexity or tau PET images, was used as input to a CNN classifier, in a 3D format with dimensions of 79×95×68. The architecture of CNN model is illustrated in Figure 2. The CNN model used in this study was composed of a series of feature maps across seven convolutional layers, structured as: 32-32-64-64-128-128-256. Between the convolutional layers, three max-pooling layers were inserted between layers 32-64, 64-128, and 128-256. A global average pooling layer was inserted before the last layer, which was a fully connected layer with a dropout ratio of 0.5 followed by a softmax function. Each convolutional layer contained a two-step process: a 3×3×3 kernel convolution followed by EvoNorm-S0.[14] EvoNorm-S0 is the combination of a normalization function and an activation function. In particular, this function showed superior performance compared to alternative methods such as batch normalization-Rectified Linear Unit (ReLU) or group normalization-ReLU. Crucially, EvoNorm-S0 consistently outperformed alternatives across various mini-batch sizes. Its robust performance on small mini-batches is crucial for 3D image training, where memory constraints limit the mini-batch size.

**Figure 1.**
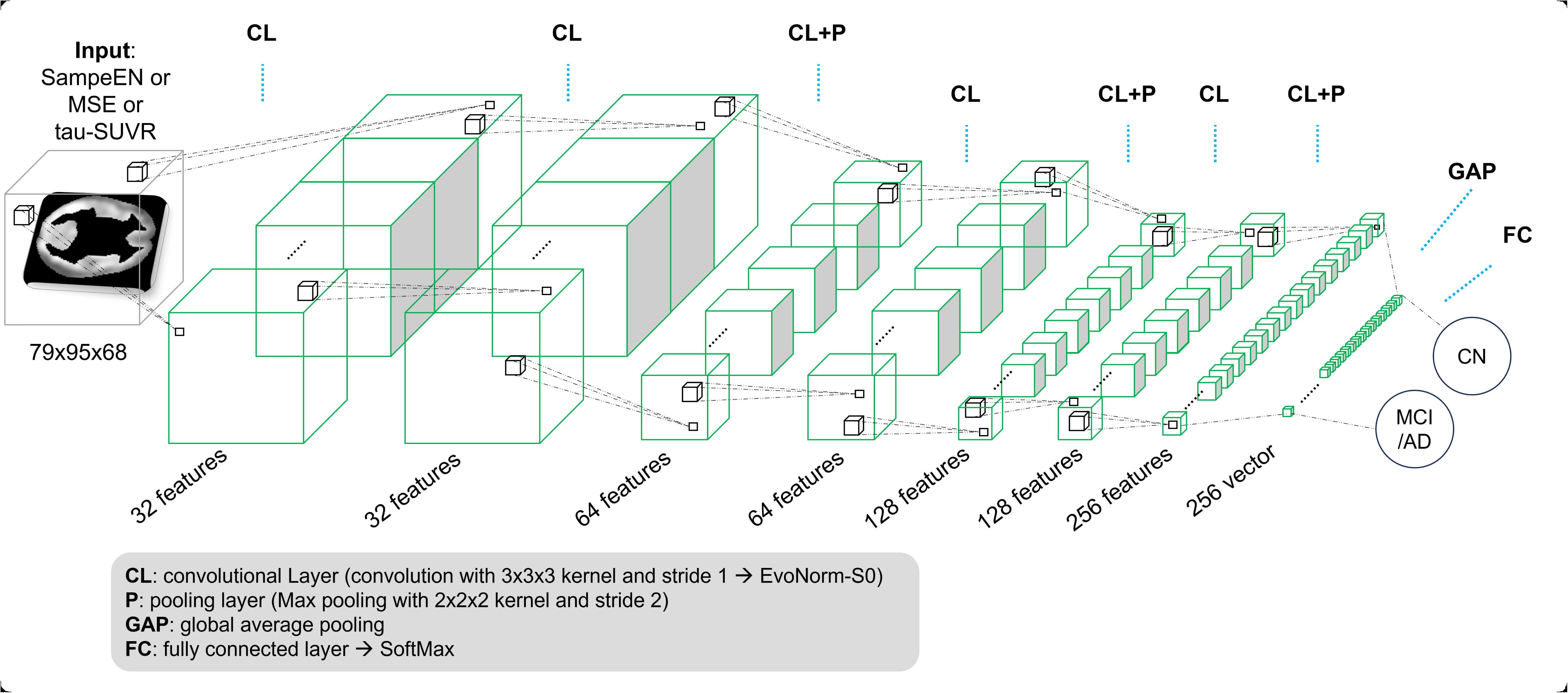
Functional Brain Networks Used for Classification. The nine functional brain networks utilized in this study, mapped onto the cortical gray matter in MNI space. These networks are based on an extended version of the seven-network parcellation defined by Yeo et al. [11], including the sensorimotor, frontoparietal, dorsal attention, ventral attention with language, default mode, visual, and limbic system networks. Additionally, the salience and auditory networks were defined to create a total of nine regions of interest. Each network is color-coded and overlaid on a standard brain surface template, illustrating the specific regions analyzed for their impact on classification performance between cognitively normal (CN) and MCI/AD subjects.

**Figure 2.**
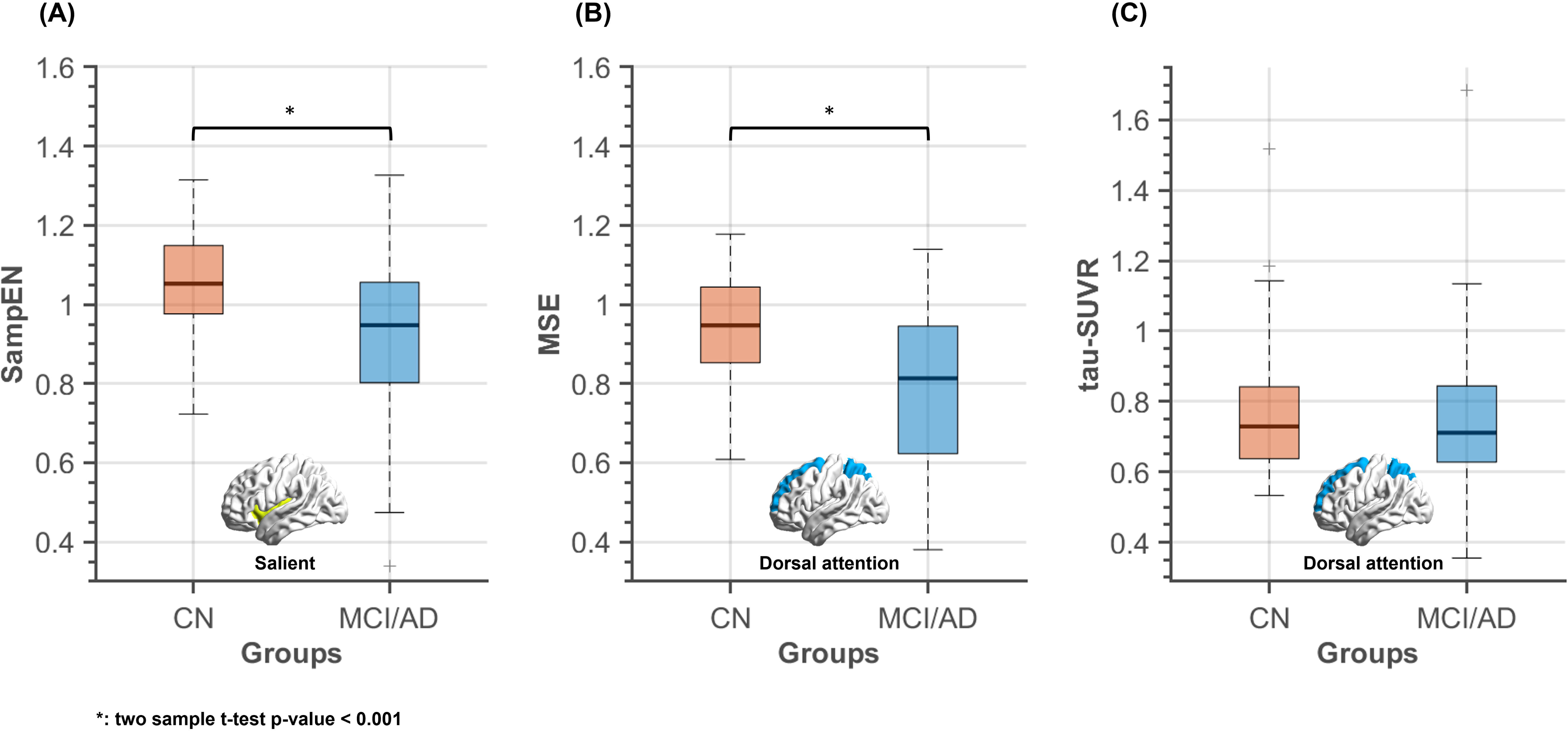
Workflow and Architecture of the 3D Convolutional Neural Network (CNN) The overall workflow for classifying subjects into cognitively normal (CN) or MCI/AD groups using either fMRI complexity maps (Sample Entropy [SampEn] or Multi-Scale Entropy [MSE]) or tau-PET SUVR images. Each image type underwent preprocessing, cortical gray matter masking, and was input into the CNN. The input layer accepts 3D images with dimensions of 79 × 95 × 68 voxels. The network consists of seven convolutional layers with feature map sizes of 32-32-64-64-128-128-256. Three max-pooling layers are placed after the third, fifth, and seventh convolutional layers. Each convolutional layer applies 1 a 3 × 3 × 3 kernel followed by the EvoNorm-S0 activation function. A global average pooling layer reduces the spatial dimensions before a fully connected layer with a dropout rate of 0.5. The final layer uses a softmax function to output the classification probability for CN or MCI/AD.

During the training phase, we performed online data augmentation through affine transformations, including scaling, translation, rotation, and shearing. A mini-batch size of 10 and a total of 300 epochs were set, with early stopping using the validation data set to mitigate overfitting. Adam was used as the optimizer with a learning rate of 0.00005. For a comprehensive evaluation, we performed a 5-fold cross-validation on the training data set and extracted the validation results. The performance evaluation metrics included accuracy, F1-score, area under the curve (AUC), precision, and sensitivity calculated for each model. Our dataset had an unbalanced class distribution, with a higher number of CN subjects compared to MCI and AD subjects. In this case, the F1-score is a particularly valuable metric for evaluating model performance. Unlike accuracy, which can be biased by the majority class in imbalanced datasets, the F1-score is computed by harmonizing precision and sensitivity. A high F1-score indicates that the model achieves a good trade-off between these two aspects, making it a robust indicator of performance even when class distributions are uneven. TensorFlow was used for the implementation to train and test the CNN models.

Each classifier was trained separately with either fMRI complexity images or SUVR images. These two models used the same network structures and hyperparameters.

### 2.6 Analyze the impact of each ROI on MCI/AD classification

To assess the individual impact of each ROI within fMRI complexity and tau-PET imaging on MCI/AD classification, we recomputed the CNN several times using a systematic leave-one ROI-out strategy. Voxels belonging to a given network ROI were zero-padded and excluded from the whole GM region, and the resulting truncated data were used as new input to the model of each modality trained previously in Section 2.5. By comparing the classification performance (accuracy, F1-score, and AUC) between the truncated input and the input using the whole GM region, we quantified the influence of the given network ROI on CN vs MCI/AD classification. We then repeated this process across 9 network ROIs. This approach allowed us to identify the most influential brain network ROI in discriminating between CN and MCI/AD subjects.

Furthermore, we extracted the mean value of fMRI complexity or Tau SUVR per subject from the most influential network ROI and examined the group difference between CN and MCI/AD. This analysis provided insight into how much the network ROI, which most contributed to classification performance, displayed differences in the two imaging feature values across the different diagnostic groups.

## 3. Results

### 3.1 Classification performance comparison

The results of the 5-fold cross-validation showed that the model with fMRI complexity generally achieved comparable or superior classification performance to the model based on tau- PET (accuracy: 0.8000; F1-score: 0.6111; precision: 0.8148; sensitivity: 0.4889; AUC: 0.6654, Table 1). Notably, the model with SampEN showed the best values for all performance metrics (accuracy: 0.8429; F1-score: 0.6944; precision: 0.9259; sensitivity: 0.5556; AUC: 0.7623), outperforming the other modalities. The model with MSE also showed better performance than tau-PET (accuracy: 0.8000; F1-score: 0.6410; precision: 0.7576; sensitivity: 0.5556; AUC: 0.7322), except for precision (worse) and accuracy (same).

**Table 1.** Classification Performance Metrics of 3D CNN Models for fMRI-Complexity and tau-PET based on whole cortical gray matter input.

| Validation |  | Accuracy | F1-score | Precision | Sensitivity | AUC |
| --- | --- | --- | --- | --- | --- | --- |
| fMRI<br>Complexity | SampEN | **0.8429** | **0.6944** | **0.9259** | **0.5556** | **0.7623** |
|  | MSE | 0.8000 | 0.6410 | 0.7576 | **0.5556** | 0.7322 |
| tauPET |  | 0.8000 | 0.6111 | 0.8148 | 0.4889 | 0.6654 |

\*Abbreviations: SampEn=Sample Entropy, MSE=Multiscale Entropy, and AUC=Area under curve

### 3.2 Influence of each network ROI on classification performance

By subtracting a classification performance metric (accuracy, F1-score, and AUC) of the whole GM region as an input from that of the input excluding a given network ROI, we evaluated the influence of the given network ROI (Table 2). A value of 0 for each metric indicates that there has been no change in performance upon exclusion of the target network ROI. Conversely, a value of -1 signifies a 100% performance drop when excluding the ROI. Models trained with SampEN consistently showed the largest drop in performance across all metrics when the salience network region was removed (accuracy: -0.1391; F1-score: -0.3277; AUC: -0.0639), while models trained with MSE were most affected by the exclusion of the dorsal attention network region (accuracy: -0.1056; F1-score: -0.3742; AUC: -0.0695). For tau-PET, the removal of the dorsal attention region led to the largest decrease in accuracy (-0.0192) and F1-score (-0.0699), while the removal of the sensorimotor network region had the largest impact on AUC (-0.0063). In conclusion, these results suggest that the dorsal attention network is a critical region for classification in both MSE and tau-PET, while the salience network is particularly important for MSE.

**Table 2:** Classification performance of 3D CNN for fMRI-Complexity and tau-PET, respectively, using truncated input data after systematically removing areas belonging to a specific network.

| Target network | Accuracy |  |  | F1-score |  |  | AUC |  |  |
| --- | --- | --- | --- | --- | --- | --- | --- | --- | --- |
|  | SampEN | MSE | tauPET | SampEN | MSE | tauPET | SampEN | MSE | tauPET |
| Sensorimotor | -0.0411 | -0.0325 | -0.0108 | -0.0871 | -0.1198 | -0.0394 | -0.0122 | -0.0070 | **-0.0063** |
| Frontoparietal | -0.0593 | -0.0268 | -0.0179 | -0.1168 | -0.0847 | -0.0455 | -0.0365 | 0.0137 | 0.0000 |
| Dorsal attention | -0.0730 | **-0.1056** | **-0.0192** | -0.1720 | **-0.3742** | **-0.0699** | -0.0240 | **-0.0695** | 0.0081 |
| Ventral with language | -0.0425 | -0.0358 | -0.0179 | -0.0521 | -0.1332 | -0.0650 | 0.0341 | -0.0071 | 0.0018 |
| Default mode | -0.0408 | -0.0472 | -0.0172 | -0.0725 | -0.1808 | -0.0537 | -0.0001 | -0.0444 | 0.0024 |
| Salient | **-0.1391** | -0.0293 | -0.0145 | **-0.3277** | -0.0913 | -0.0526 | **-0.0639** | -0.0038 | 0.0124 |
| Auditory | -0.0849 | -0.0358 | -0.0178 | -0.2151 | -0.1114 | -0.0641 | 0.0645 | -0.0371 | 0.0056 |
| Visual | -0.0779 | -0.0315 | -0.0189 | -0.0887 | -0.1014 | -0.0272 | -0.0312 | -0.0365 | 0.0022 |
| Limbic system | -0.0545 | -0.0063 | -0.0128 | -0.1108 | -0.0196 | -0.0464 | -0.0210 | 0.0014 | 0.0082 |

\*Abbreviations: SampEn=Sample Entropy, MSE=Multiscale Entropy, and AUC=Area under curve

Furthermore, we computed the mean value of the most influential network ROI for each neuroimaging approach (salience network for SampEn; dorsal attention network for MSE and tau-PET) and compared them between CN and MCI/AD groups. As shown in Figure 3, the SampEn values within the salience network were significantly higher in the CN group compared to the MCI/AD group (mean ± std: 1.0530 ± 0.1233 vs. 0.9457 ± 0.1102; p < 0.001). Similarly, the MSE values within the dorsal attention network was significantly higher in the CN group compared to the MCI/AD group (mean ± std: 0.9399 ± 0.1279 for CN vs. 0.7767 ± 0.2008 for MCI/AD; p < 0.001). The tau-PET SUVR values within the dorsal attention network were slightly, but not significantly higher in the MCI/AD group than in the CN group (mean ± std: 0.7595 ± 0.2371 vs. 0.7518 ± 0.1577).

**Figure 3.**
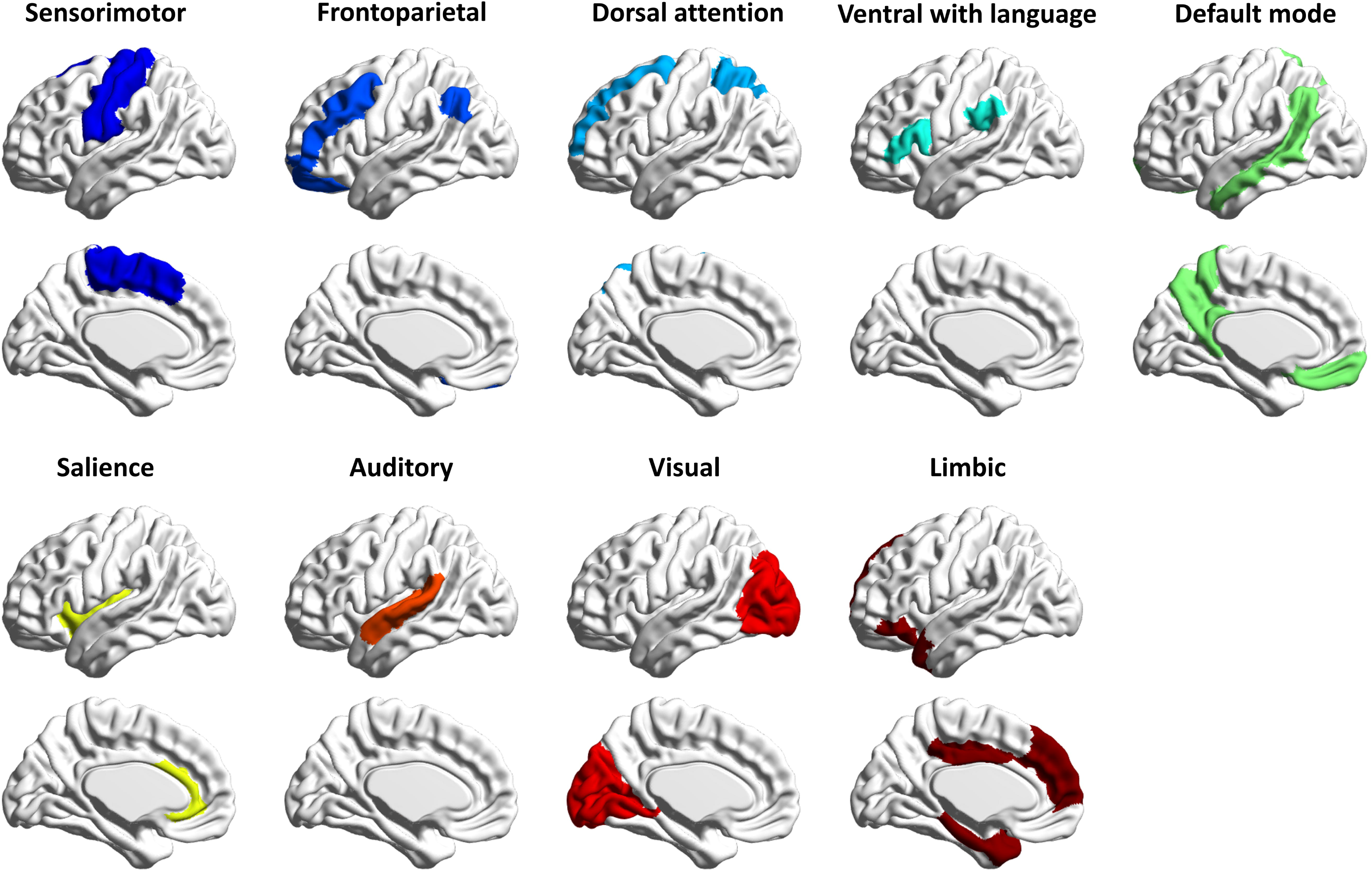
Imaging Value Comparisons Within the Most Influential Networks for Classification. Comparison of imaging values between cognitively normal (CN) and MCI/AD groups within the most influential brain networks identified for each imaging modality. **(A)** Mean Sample Entropy (SampEn) values within the salience network, showing significantly higher complexity in the CN group compared to the MCI/AD group (mean ± SD: CN 1.0530 ± 0.1233 vs. MCI/AD 0.9457 ± 0.1102; **p** < 0.001). **(B)** Mean Multi-Scale Entropy (MSE) values within the dorsal attention network, also significantly higher in the CN group (mean ± SD: CN 0.9399 ± 0.1279 vs. MCI/AD 0.7767 ± 0.2008; **p** < 0.001). **(C)** Mean tau-PET SUVR values within the dorsal attention network, with no significant difference between groups (mean ± SD: CN 0.7518 ± 0.1577 vs. MCI/AD 0.7595 ± 0.2371). These findings highlight the sensitivity of fMRI complexity measures in detecting group differences within specific networks associated with cognitive function.

## 4. Discussion

The objective of this study was to evaluate whether fMRI-complexity can serve as an additional neuroimaging marker to classify subjects with cognitive impairment in the Alzheimer’s Disease (AD) spectrum from those with normal cognition. We hypothesized that fMRI-complexity reflects the brain’s capacity of information processing and cognitive flexibility and correlates with cognitive function. Thus, we tested whether classification performance of fMRI-complexity is comparable to the state-of-the-art method using tauPET. Our study employed the two most used fMRI-complexity measures, Sample Entropy (SampEn) and its temporal expansion Multiscale Sample Entropy (MSE). Our classification results using a 3D- CNN model suggest that both complexity measures can predict cognitive impairment in AD spectrum with accuracy comparable to tauPET.

Most prior studies, like ours, have utilized the ADNI dataset,[15] primarily focusing on classification between cognitively normal (CN) and AD due to the larger cognitive and neurophysiological differences, which facilitate classification. Fewer studies have explored CN vs. mild cognitive impairment (MCI),[16] and even fewer have used fMRI for classification, as PET and structural MRI are more common.[16] Our study addresses this gap by using fMRI complexity for classification between cognitively impaired and unimpaired individuals. We achieved an accuracy of around 85%, consistent with prior machine learning approaches using fMRI.[17] Notably, while MSE classification showed reduced complexity in the Dorsal Attention Network (DAN), similar to the pattern seen in tauPET-based classification, SampEn classification indicated reduced complexity in the Salience Network (SAL). This suggests that MSE and SampEn capture distinct pathophysiological processes. High-frequency fluctuations detected by SampEn may reflect local neuronal changes, while the slower frequencies of MSE could reflect integrative processing in distributed networks (Wang 2018). Supporting this, SampEn shows little or negative correlation with functional connectivity (FC), while MSE correlates positively with FC.[18]

Both, SampEn and MSE, have been demonstrated to show spatially defined alterations among CN subjects, patients with MCI and those with AD. While findings for SampEn can vary across studies —potentially due to differences in TR and the frequencies captured by SampEn — MSE results have been more consistent, typically showing a decline in complexity with increasing cognitive impairment in AD progression. We recently showed that MSE is negatively associated with tauPET, not only in the medial temporal lobe but also in areas corresponding to the DAN such as the lateral prefrontal and parietal cortex. Furthermore, decreased fMRI complexity in lateral prefrontal cortex and lateral parietal lobe is associated with impaired cognitive function.[6] Notably, fMRI complexity appears to be specifically related to tau protein deposition but not amyloid-beta deposition,[19] thus corroborating the assumption that fMRI complexity is sensitive to tau-related disruptions in neuronal signaling, which subsequently contribute to cognitive impairment. In summary, MSE and tau PET appear to have neurophysiological and neurocognitive interrelations while SampEn’s underlying neurophysiological mechanisms are less defined. Nevertheless, SampEn in our study achieved comparable classification performance, highlighting its association with the SAL, another key network involved in cognitive control and allocation of cognitive resources. Dysfunction of the SAL has been linked to AD,[20] likely due to its role in reallocating resources between internal (DMN) and external (DAN) attentional processes,[21] a cognitive function that shows deficits in AD.[22] Thus, SampEn might be sensitive to early local changes in allocation attentional processes, while MSE is sensitive changes in distributed information processing, both critical aspects of cognitive impairments in the AD progression.

While our study has methodological strengths, we acknowledge certain limitations. The classification performance using tau-PET in our study was lower than reported in prior studies.[23,24] This may be due to our relatively small sample size, as only a limited subset of ADNI participants had both fMRI and tau-PET data, limiting the data available for model training. Additionally, to ensure comparability with fMRI complexity, we restricted our analysis on the cerebral cortex rather than the entire brain (no white matter or no deep gray matter structures included). This focus may have reduced the overall performance of the tau-PET-based classification. However, since our primary objective was to compare the performance of fMRI complexity and tau-PET under identical conditions using the same deep neural network architecture and preprocessing procedure, we believe these limitations do not undermine our findings. Our results provide valuable insights into the potential of fMRI complexity as a surrogate biomarker for tau-PET in AD classification.

In summary, the classification performance suggests that fMRI-MSE could serve as a surrogate biomarker to tau-PET for AD. While fMRI requires more time for acquisition, preprocessing, noise reduction, and analysis compared to structural MRI, it provides a unique view of functional changes that may precede structural alterations. Unlike PET, fMRI is non- invasive, and specifically, fMRI complexity has been proposed to be sensitive to tau-related neuronal disruption and resultant brain function. Thus, fMRI-complexity-based classification may be particularly valuable for detecting early-stage brain physiological changes and distinguishing cognitively normal individuals from those with cognitive impairment. However, further studies using larger, multisite datasets are needed to validate these preliminary findings and strengthen the evidence.

## Acknowledgements

Data collection and sharing for the Alzheimer’s Disease Neuroimaging Initiative (ADNI) is funded by the National Institute on Aging (National Institutes of Health Grant U19 AG024904). The grantee organization is the Northern California Institute for Research and Education.

In the past, ADNI has also received funding from the National Institute of Biomedical Imaging and Bioengineering, the Canadian Institutes of Health Research, and private sector contributions through the Foundation for the National Institutes of Health (FNIH) including generous contributions from the following: AbbVie, Alzheimer’s Association; Alzheimer’s Drug Discovery Foundation; Araclon Biotech; BioClinica, Inc.; Biogen; Bristol-Myers Squibb Company; CereSpir, Inc.; Cogstate; Eisai Inc.; Elan Pharmaceuticals, Inc.; Eli Lilly and Company; EuroImmun; F. Hoffmann-La Roche Ltd and its affiliated company Genentech, Inc.; Fujirebio; GE Healthcare; IXICO Ltd.; Janssen Alzheimer Immunotherapy Research & Development, LLC.; Johnson & Johnson Pharmaceutical Research & Development LLC.; Lumosity; Lundbeck; Merck & Co., Inc.; Meso Scale Diagnostics, LLC.; NeuroRx Research; Neurotrack Technologies; Novartis Pharmaceuticals Corporation; Pfizer Inc.; Piramal Imaging; Servier; Takeda Pharmaceutical Company; and Transition Therapeutics.

